# SwarmGenomics: A Unified Pipeline for Individual-Based Whole-Genome Analyses

**DOI:** 10.1101/2025.08.13.670070

**Authors:** Aure Kylmänen, Yu-Chi Chen, Sahar Javaheri Tehrani, Nikolas Vellnow, Justin J. S. Wilcox, Toni I. Gossmann

**Affiliations:** TU Dortmund University, Computational Systems Biology, Faculty of Biochemical and Chemical Engineering, Emil-Figge-Str. 66, 44227 Dortmund, Germany

**Keywords:** Whole-genome sequencing, Bioinformatics pipeline, Heterozygosity, Runs of homozygosity, PSMC, NUMT, Mitochondrial genome, Repeat annotation, Unmapped reads

## Abstract

Advances in sequencing technologies have made whole-genome data widely accessible, enabling research in population genetics, evolutionary biology, and conservation. However, analyzing whole-genome sequencing (WGS) data remains challenging, often requiring multiple specialized tools and substantial bioinformatics expertise. We present SwarmGenomics, a modular, user-friendly command-line pipeline for reference-based genome assembly and individual-based genetic analyses. The pipeline integrates seven modules: heterozygosity estimation, runs of homozygosity detection, Pairwise Sequentially Markovian Coalescent (PSMC) analysis, unmapped reads classification, repeat analysis, mitochondrial genome assembly, and nuclear mitochondrial DNA segment (NUMT) identification. Each module can be run independently or as part of a complete workflow. We demonstrate the pipeline’s utility with a case study on the giant panda (*Ailuropoda melanoleuca*), revealing insights into genetic diversity, inbreeding history, historical population size changes, transposable element activity, and microbial contamination. SwarmGenomics lowers the entry barrier for genomic analysis of diploid, non-model species, serving both as a research and teaching tool. The pipeline and documentation are available at https://github.com/AureKylmanen/Swarmgenomics.

## Motivation

Whole-genome sequencing (WGS) data is increasingly accessible due to significant advancements and declining costs of sequencing technologies. A primary catalyst for this development was the Human Genome Project, which spurred innovations in high-throughput sequencing and bioinformatics (Hood and Rowen 2013). This project also led to the establishment of publicly accessible genomic databases, such as GenBank, Ensembl, and the UCSC Genome Browser (Birney et al. 2009). The rapid accumulation of genomic information in these databases has begun to outpace the availability of computational resources and analytical tools necessary for its effective use. Complementary initiatives, such as Bird10K (Prum et al. 2015) and Bat1K (Teeling et al. 2018), have further contributed to the expansion of genomic resources. Currently, the NCBI genome database contains genomic data for approximately 4,264 mammalian, 2,096 avian, and 2,578 reptilian species (NCBI 2025a), representing a continually growing resource that supports research across more than 6,400 mammalian species and approximately 11,000 bird and reptile species. This expanding genomic resource creates unprecedented opportunities for comparative and evolutionary studies across a wide range of species.

In parallel, large-scale genome consortia such as the Vertebrate Genomes Project (Rhie et al. 2021) and the Earth BioGenome Project (Lewin et al. 2022) are generating large volumes of high-quality reference genomes. As these comprehensive reference genomes become available, there is an increasing emphasis on moving beyond just genome assembly towards in-depth population genomics analyses. Such analyses are crucial for uncovering patterns of genetic variation within and between populations, identifying signatures of natural selection, and better understanding the evolutionary history of species. Moreover, detailed population genomic data are essential for informing conservation strategies, particularly for endangered species facing threats from habitat loss, climate change, and human activity. As the number of reference genomes continues to grow along with improvements in sequencing technologies, the demand for accessible and reliable tools is becoming more pressing.

Despite significant progress, the analysis of whole-genome sequencing data remains technically and computationally demanding, often requiring extensive bioinformatics and programming expertise. This presents a substantial barrier for researchers from diverse scientific backgrounds. Laboratories are sometimes required to hire dedicated bioinformatics staff to overcome these challenges (Oakeson et al. 2017). The typical genomic analysis workflow involves multiple specialized software packages and several programming languages, leading to fragmented processes that can be especially daunting for beginners. To date, integrated pipelines that simplify and streamline genomic analyses into an accessible workflow remain limited.

Some tools such as the web-based platform Galaxy (Afgan et al. 2016) and pipelines like Genome After Party (Sanger 2025) have made significant progress lowering the entry barrier to bioinformatics by offering more userfriendly, integrated environments for genomic analyses. While Galaxy provides a graphical interface for many popular tools, Genome After Party focuses on streamlined, reproducible workflows for genome assembly and annotation. However, handling large whole-genome datasets or customising complex workflows may still require bioinformatics expertise. Furthermore, these workflows and pipelines are not specifically optimised for population genomics or the analysis of diploid, non-model organisms.

To address these limitations, we developed SwarmGenomics, an integrated, user-friendly bioinformatics pipeline designed to facilitate reference-based genome assembly and genetic analyses. Specifically optimised for diploid, non-model organisms, SwarmGenomics broadens the accessibility of genomic analyses for species that remain understudied. The pipeline encompasses a straightforward, step-by-step assembly procedure followed by several analytical modules, including heterozygosity analysis, detection of runs of homozygosity, Pairwise Sequentially Markovian Coalescent (PSMC) analysis, characterization of unmapped reads, repeat element annotation, mitochondrial genome assembly, and identification of nuclear mitochondrial DNA segments (NUMTs). SwarmGenomics is particularly well-suited for researchers with limited bioinformatics training, and is intended to serve as both an educational resource and a self-learning tool.

To illustrate the practicality and educational value of SwarmGenomics, we demonstrate its application using real genomic data from the giant panda (*Ailuropoda melanoleuca*), an iconic species of conservation interest. This case study highlights the pipeline’s effectiveness as an accessible entry point into genomic research, providing insights into population genetics, evolutionary biology, and conservation genetics.

## Pipeline and Pre-processing

SwarmGenomics is a versatile and user-friendly bioinformatics pipeline developed for whole-genome sequencing (WGS) analysis starting directly from raw paired-end reads by default, but can be adapted to single-end data as well. An overview of the pipeline’s workflow is shown in Figure 1, with prospective future modules indicated in grey. Its modular design enables users to select and execute only the analyses relevant to their specific research needs (Table 1). The pipeline is implemented primarily in Bash, Python, and R; however, no prior experience with these programming languages is necessary to use it effectively. We recommend running SwarmGenomics within a virtual machine (VM) to simplify software setup and enhance reproducibility in genomic analyses (Nocq et al. 2013). Alternatively, users without access to a VM can run the pipeline on a dedicated Linux workstation or a sufficiently powerful laptop.

**Table 1:**
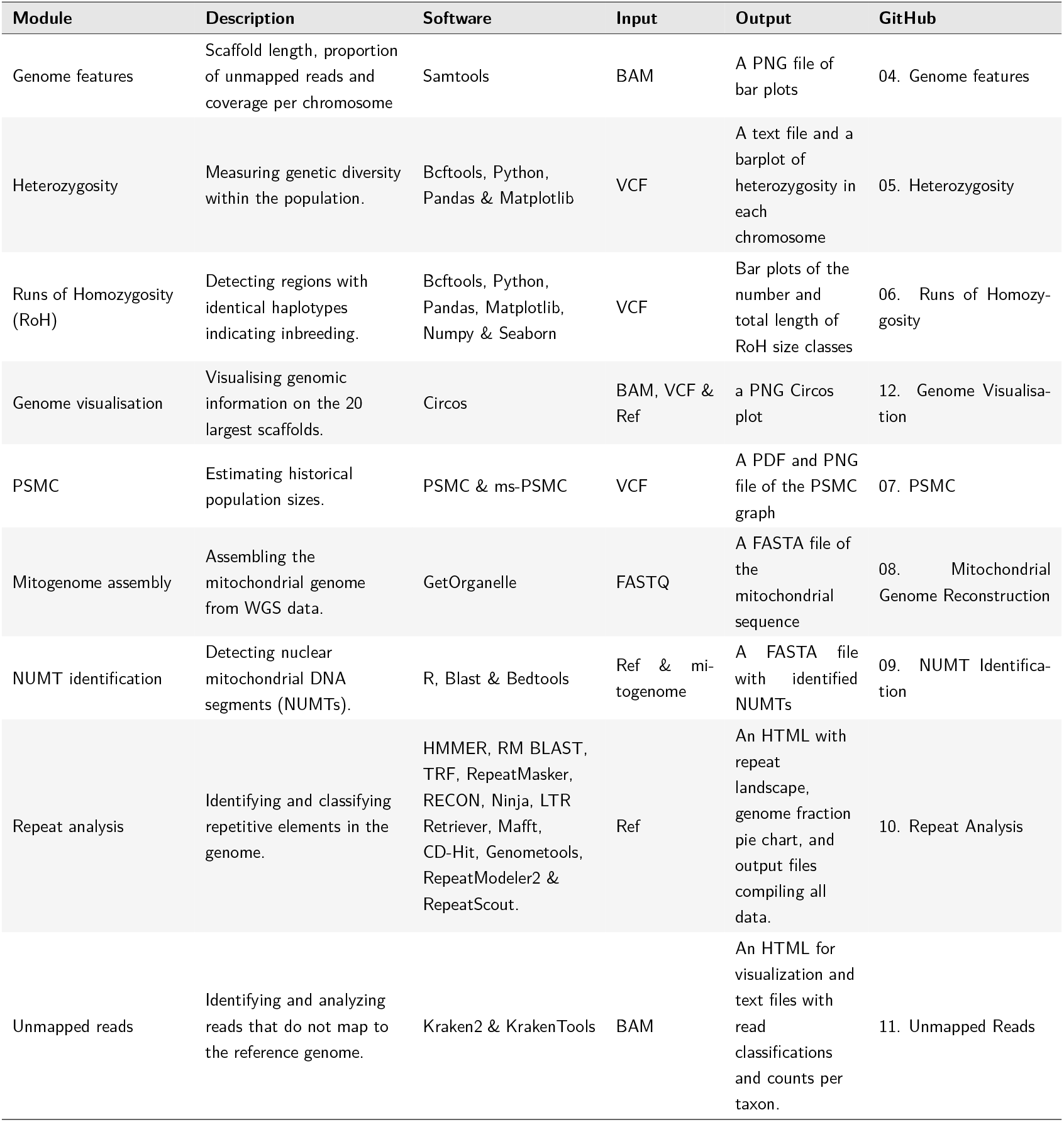
Genomic analysis modules in SwarmGenomics. Overview of each module’s description, required software, input, output format, and source code availability

**Figure 1:**
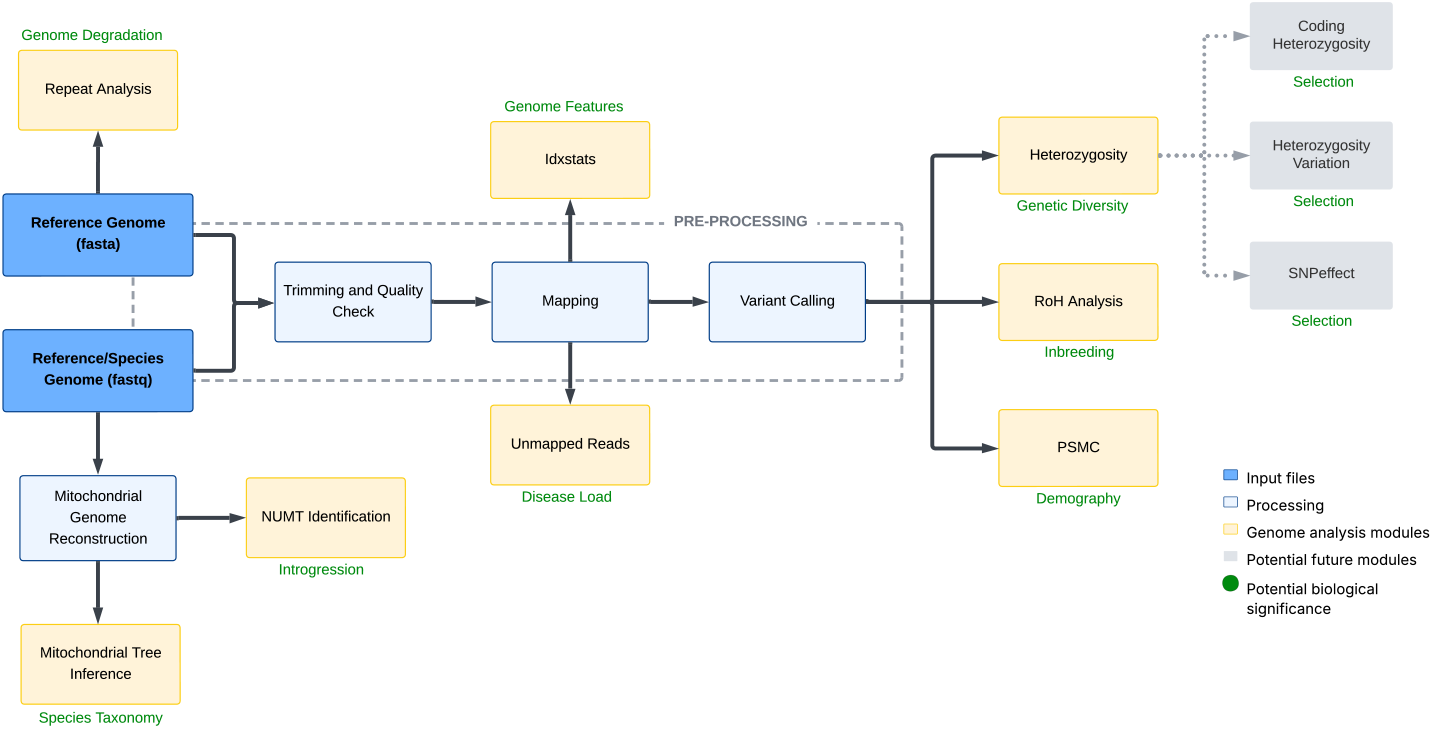
Overview of the SwarmGenomics workflow. SwarmGenomics pipeline overview starting with reference and species genomes in FASTA and FASTQ format, followed by processing steps (in light blue) and modules for different genomic analyses (in yellow), with possible future modules in grey.

### Mapping

| (Virtual) Machine | Data, Software and Time Requirements |
| --- | --- |
| 8-20 (V)CPUs<br>+16 GB RAM<br>300–400 GB storage | **Input:** Reference genome (FASTA format) and reads from sequenced individual (FASTQ format)<br>**Installations:** Miniconda3, FastQC, bwa-mem2, samtools, trimmomatic, SRA Tools<br>**Time:** <5h |

SwarmGenomics is designed to leverage the diploid genome of a single individual in combination with a reference genome, to infer population genetic characteristics at the species or population level. The input genomic data (i.e. FASTQ files) may or may not originate from the same individual as the reference genome. It is important that datasets are clearly labelled to indicate whether they represent a reference individual or another sample from the same or a related species. For most modules, two genomic inputs are required: (1) a reference genome, ideally high-quality and species-specific, and (2) raw sequencing reads (in FASTQ or SRA format) from a single diploid individual. These can be derived from the same individual, although this is not strictly necessary. SwarmGenomics can be used to assess how well the input data match, and whether the individual aligns closely with the selected reference genome.

The reference genome is typically a well-annotated, curated sequence that represents the consensus DNA of a species, whereas the individual genome reflects the unique genetic composition of the sequenced sample. Both genome types are accessible through public databases such as NCBI Genomes and the Sequence Read Archive (SRA) (NCBI 2025a; NCBI 2025b), as well as other repositories like ENA, DDBJ, or species-specific consortia (O’Cathail et al. 2025; Tanizawa et al. 2023; Zoonomia Consortium 2020; Smith et al. 2023). It is recommended to use a reference genome from RefSeq, as these assemblies are curated and annotated for accuracy (O’Leary et al. 2016). For the sequencing reads, whole-genome, paired-end Illumina short reads from a representative individual of the species (or a closely related species) are preferred, but in principle long read data from PacBIO or Oxford Nanopore (Eid et al. 2009; Jain et al. 2016) may be used although the pipeline has not been specifically developed for these.

Prior to downstream genetic analyses, raw sequencing data must be pre-processed and assembled. Genome assembly is a foundational step that reconstructs the genome from sequencing reads and enables the identification of variants and genomic features. While reference-based genome assembly can be complex, SwarmGenomics simplifies this process through a guided, step-by-step protocol and integrated preprocessing scripts.

Instructions for installing the required software are provided on the SwarmGenomics GitHub page (github.com/AureKylmanen/Swarmgenomics). For users starting with SRA files, these are first converted to FASTQ format, the standard format for storing raw sequencing reads and their associated quality scores (Cock et al. 2010). Quality control is performed using FastQC (Andrews et al. 2010), which assesses read quality, base composition, and detects overrepresented sequences. Trimming is carried out with Trimmomatic (Bolger et al. 2014) to remove low-quality bases and adapter contamination. A second FastQC check is recommended to verify improvements in read quality.

Subsequently, reads are aligned to the reference genome using BWA (H. Li and Durbin 2009). The reference genome must first be indexed with `bwa index`. Paired-end reads are then aligned using `bwa mem`, producing a SAM file. This file is sorted and converted to BAM format using `samtools sort`, and then indexed with `samtools index` (H. Li, B. Handsaker, et al. 2009). The reference genome is also indexed with `samtools faidx` to facilitate random access during variant calling.

### Variant calling

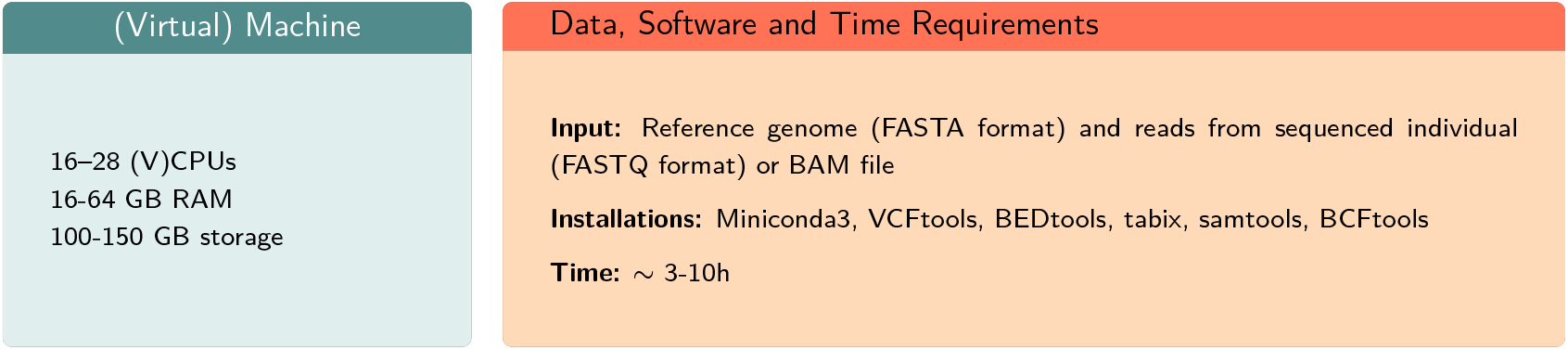

Variant calling is a fundamental step in identifying genomic differences between an individual’s genome and the reference. In SwarmGenomics, this is performed using `bcftools mpileup`, a lightweight and widely used tool that efficiently computes genotype likelihoods and generates a VCF file containing base-level information and variant calls across the genome (Danecek et al. 2011). BCFtools is used for its speed, simplicity, and compatibility with standard short-read alignments, making it a practical default option for most users. The resulting VCF file serves as a key input for multiple downstream modules within the pipeline.

For users interested in more sophisticated or fine-tuned variant calling approaches, several alternative tools are available. FreeBayes is a haplotype-based caller that supports multi-sample joint calling and better resolves complex variants, although it typically requires more parameter tuning. GATK HaplotypeCaller is widely regarded for its robustness and accuracy, especially in well-annotated genomes, but comes with more stringent dependencies and a steeper learning curve. ANGSD (Analysis of Next Generation Sequencing Data) is particularly useful for population-level analyses and low-coverage data, as it focuses on estimating genotype likelihoods and allele frequencies rather than making discrete genotype calls (Korneliussen et al. 2014).

While SwarmGenomics uses BCFtools for its balance of speed and usability, users with more specific needs or higher data quality may benefit from exploring these alternatives. We encourage advanced users to consider other variant callers if their analyses demand increased sensitivity, support for population-scale datasets, or improved detection of structural variants.

SwarmGenomics supports workflows that begin either from raw FASTQ files or from pre-aligned BAM files. In cases where BAM files are used, it is important that the file represents only a single diploid individual. This is especially relevant when FASTQ data come from multiple sequencing runs, each labeled with a distinct read group (RG tag). Users must ensure that all read groups correspond to the same biological individual. Details about handling multiple read groups and ensuring proper sample-level variant calling are provided in the variant calling script section on the GitHub page.

Before proceeding to downstream analyses, repetitive genomic regions are filtered from the VCF file. This is done by generating a BED file of repeat elements based on the reference genome, which is then used to mask these regions. Filtering out repeats helps to avoid false variant calls and leads to more accurate estimates of heterozygosity and other metrics derived from the variant data.

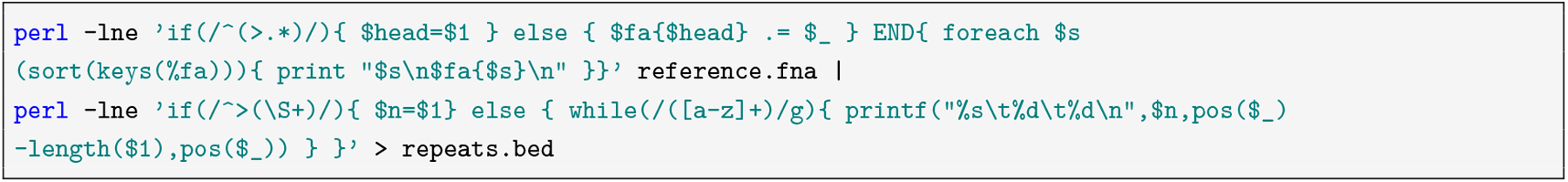

The `bedtools subtract -header` command is then used to remove the identified repeats from the VCF file. This file serves as the main input for many of the modules and analyses below. Lastly, only the largest chromosomes or scaffolds can be extracted to save time during analyses. In this example, scaffolds larger than 5 MB are extracted, though the size can be adjusted according to specific requirements.

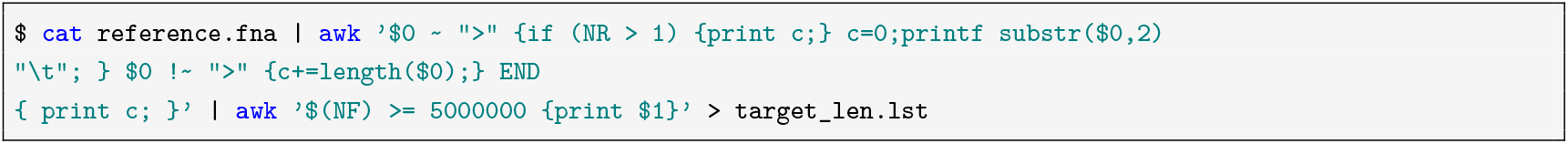

This produces a file named target_len.lst that contains the names of sequences in the reference FASTA file that are at least 5,000,000 bp long. Afterwards, these specific chromosomes are selected using `tabix`, filtered based on depth and allele count using `vcftools`, and then compressed and indexed with `bgzip` and `bcftools`. Consensus sequences in FASTQ format for each chromosome are then generated using a reference genome and the filtered VCF files, and combined into a single diploid FASTQ file, which will be used for PSMC, and the separate VCF files are used for heterozygosity and runs of homozygosity analyses. Overall the preprocessing step takes around 8-20h depending on the size of the genome and SRA files.

### List of Modules for Genome Analysis

## Genome Analysis

### Genome Features

Genome Analysis Genome Features
| (Virtual) Machine | Data, Software and Time Requirements |
| --- | --- |
| 1 (V)CPU<br>4 GB RAM<br>10 GB disk | **Input:** BAM file (or CRAM file + reference genome)<br>**Installations:** Samtools, R, ggplot2, dplyr<br>**Time:** <1 min |

When analysing genomic data, an initial overview of the mapping results is crucial for informing subsequent analyses. `Samtools idxstats` (H. Li, B. Handsaker, et al. 2009) offers a fast and efficient method for summarizing alignment statistics, providing chromosome- or scaffold-level details such as sequence length, the number of mapped reads, and the number of unmapped reads. These summary metrics can highlight patterns in sequencing coverage and mapping efficiency, and may also reveal anomalies, such as poorly assembled chromosomes or scaffolds with coverage patterns indicative of heterogametic chromosomes. In species with heteromorphic sex chromosomes (e.g., mammals), sequencing coverage can also be used to infer the biological sex of individuals by comparing read depth across sex chromosomes. Typically, males exhibit reduced or no coverage on the Y chromosome, while females lack it entirely.

These statistics can be generated using the following `samtools idxstats` command on a BAM file:

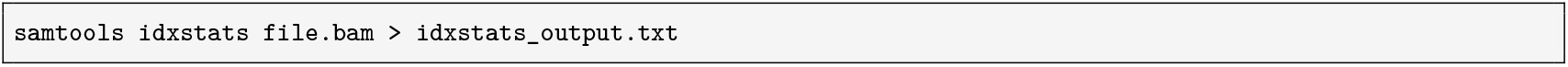

To better visualize and interpret this information, a bar plot of the 20 longest scaffolds or chromosomes (or fewer if fewer are available) is generated to highlight genomic scale. A normalized measure of unmapped reads per megabase is calculated to account for scaffold size, and an estimate of coverage per scaffold is derived based on the number of mapped reads and an assumed read length. Additionally, the total number of scaffolds and the number of scaffolds larger than 5 Mb can be obtained from the output using simple Bash commands.

The giant panda BAM file contains 22 scaffolds larger than 5 Mb. As shown in Figure 2A–C, the graphs display the size distribution of these scaffolds, the number of unmapped reads per Mbp, and the estimated coverage based on a read length of 150 bp, determined as the most frequent length among the first 1,000 alignments. The overall genome coverage is approximately 50X and remains consistent across the scaffolds.

**Figure 2:**
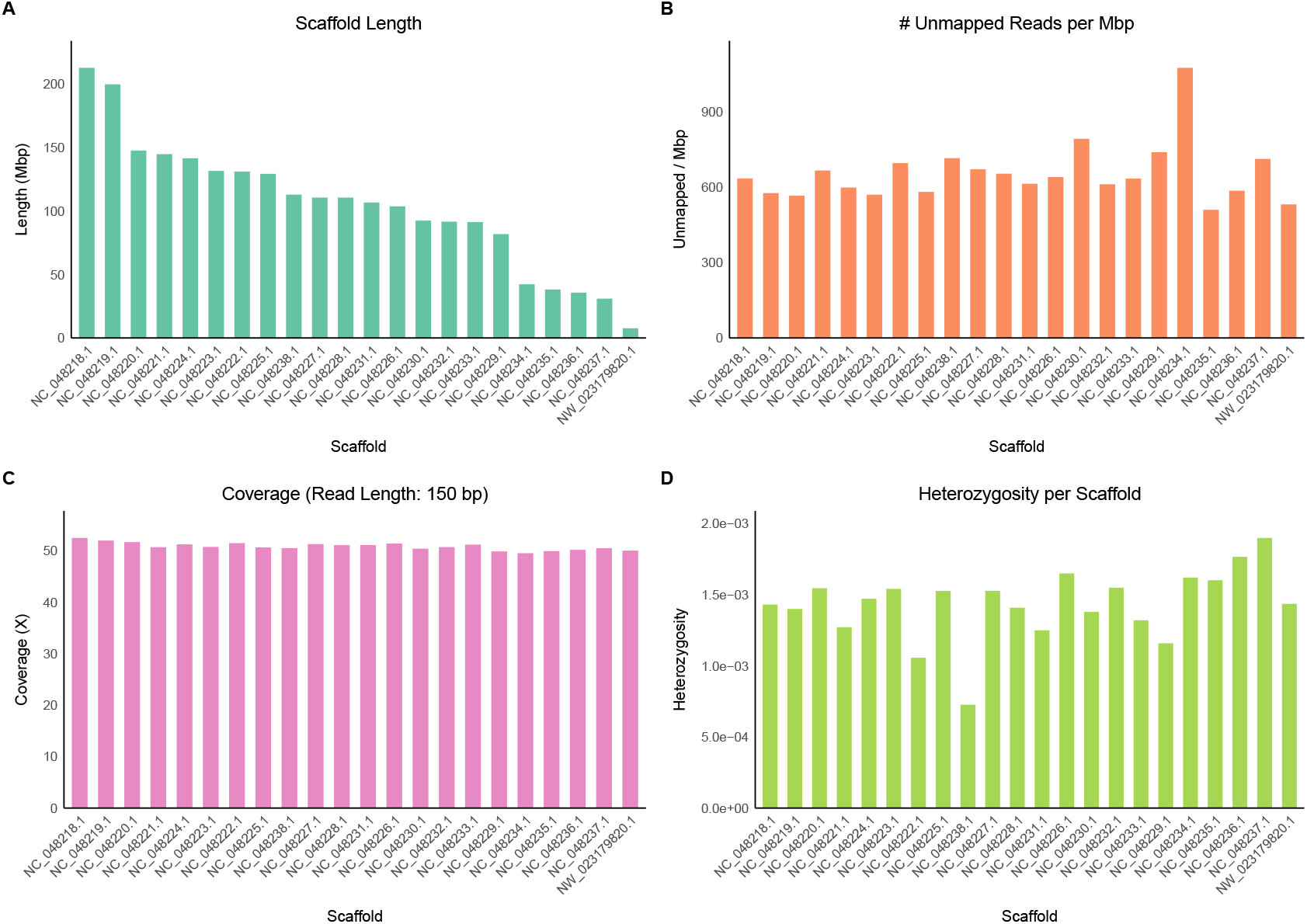
Scaffold-level genome metrics of a single individual mapped to the giant panda genome. Shown are the 22 longest scaffolds/chromosomes. **(A)** Lengths in megabasepairs (Mbp) of each scaffold. **(B)** Number of unmapped reads (i.e. paired reads missing a mate) per megabase, indicating variation in mapping efficiency across scaffolds. **(C)** Estimated sequencing coverage (X) with 150 bp read length, highlighting differences in sequencing depth among scaffolds (i.e. approximately half the coverage of the sex chromosome in an heterogametic individual). **(D)** Heterozygosity (Heterozygous sites per Mbp) of the largest scaffolds of giant panda genome

### Heterozygosity

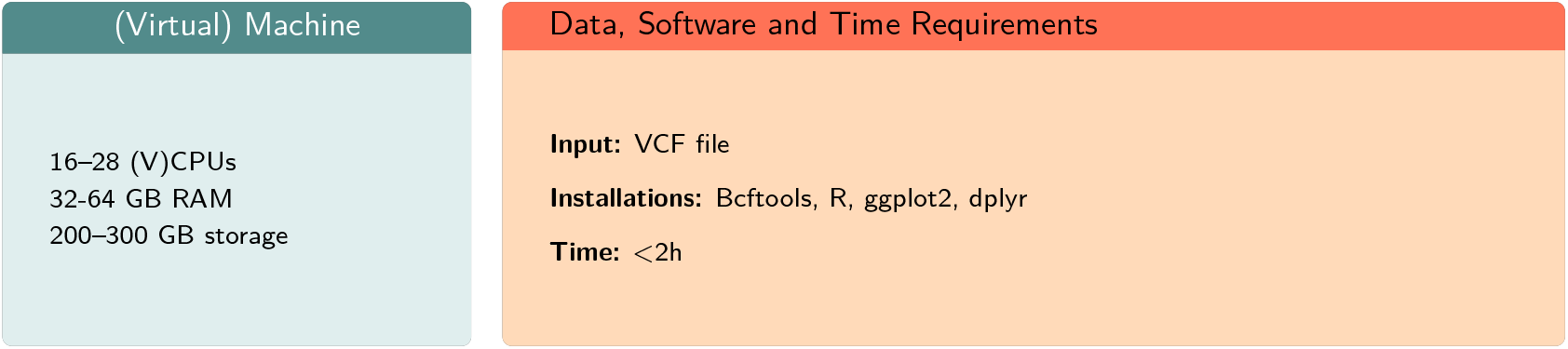

Understanding genetic diversity within a population is crucial for addressing the complexities of evolution, adaptation, and conservation. Heterozygosity, one of the most significant indicators of genetic diversity, measures the presence of different alleles at a gene locus within an individual’s genome (Nei 1973). High heterozygosity indicates healthy genetic variation, allowing populations to adapt to environmental changes and avoid inbreeding depression (Reed and Frankham 2003). Conversely, low heterozygosity may suggest inbreeding, genetic drift, or population bottlenecks, which can increase the risk of extinction (Charlesworth and Willis 2009). In the case of the giant panda, a species of conservation concern (IUCN 2023), analysing genetic diversity can provide insights into its potential to persist in a changing environment.

For estimating heterozygosity in the largest scaffolds, a bash script is used to count the number of heterozygous and homozygous genotypes, followed by calculating the proportion of heterozygous genotypes for each scaffold separately. The following Bash snippet estimates heterozygosity by counting heterozygous and homozygous genotypes from a VCF file:

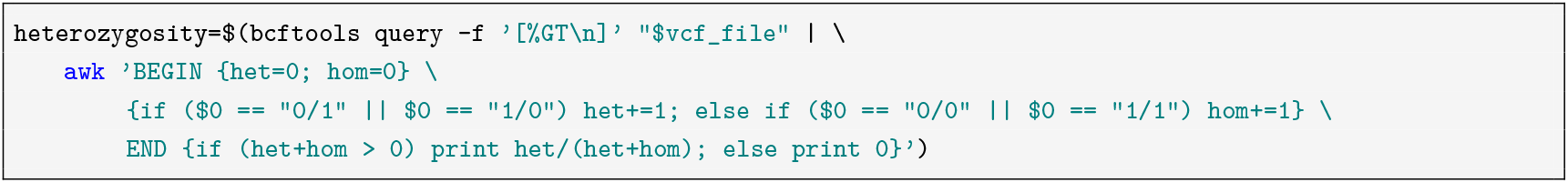

An R script can then be applied to plot the results as a bar plot, as shown in Figure 2D. Each bar represents a chromosome or a scaffold. In the case of the giant panda, heterozygosity per chromosome is moderate. However, scaffold `NC_048238.1` exhibits much lower heterozygosity than the others.

### Runs of Homozygosity

| (Virtual) Machine | Data, Software and Time Requirements |
| --- | --- |
| 16-28 (V)CPUs<br>32-64 GB RAM<br>200-300 GB storage | **Input:** VCF file<br>**Installations:** Bcftools, R; ggplot2, dplyr, gridExtra, tidyr<br>**Time:** <1h |

Similar to heterozygosity, runs of homozygosity (RoH) measure genetic diversity within populations, providing insights into genomic regions where individuals inherit identical haplotypes from both parents (Broman and Weber 1999). RoHs indicate recent common ancestry or inbreeding events, therefore the number and length of RoH can inform the demographic history of the species (Ceballos et al. 2018). High frequency and long length of RoH may suggest reduced genetic diversity and increased inbreeding depression, potentially impacting the species’ ability to adapt and survive in changing habitats (Abascal et al. 2016; Ceballos et al. 2018). Conversely, shorter or fewer RoH segments reflect greater genetic heterozygosity and potential resilience to environmental stressors. For species like the giant panda, which faces conservation challenges due to fragmented populations and vulnerability to environmental changes, RoH analysis helps assess the genetic health of the population and its resilience to environmental change.

To analyse RoH in the genome, the BCFtools roh command is used. As with heterozygosity analysis, the input file should be a VCF file with FORMAT columns containing either genotype likelihoods (PL) or genotypes (GT). By default, the command uses genotype likelihoods unless otherwise specified with the `-G` option. The command also requires an estimate of the alternate allele frequency in the population for each site; here, the default alternate allele frequency value of 0.4 is used with `--AF-dflt 0.4`.

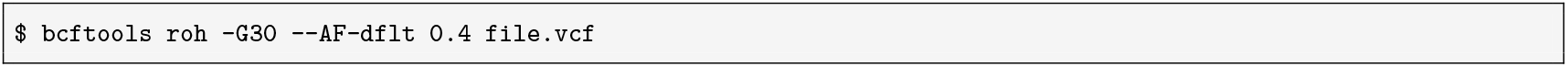

For visualizing the results, an R script is used to create bar plots of the number of RoHs and the total length of RoHs of four size categories; 10kbp – 0.1 Mbp, 0.1 - 1 Mbp, 1 - 3 Mbp, and >3 Mbp, as shown in Figure 3. Short ROHs (<0.1Mbp) result from ancient recombination and are common across populations. Short-to-medium ROHs (0.1 – 1 Mbp) indicate background relatedness within populations, while medium-to-long ROHs (1 – 3 Mbp) suggest recent shared ancestry. Long ROHs (>3Mbp) arise from recent inbreeding, where parents share a close common ancestor.

**Figure 3:**
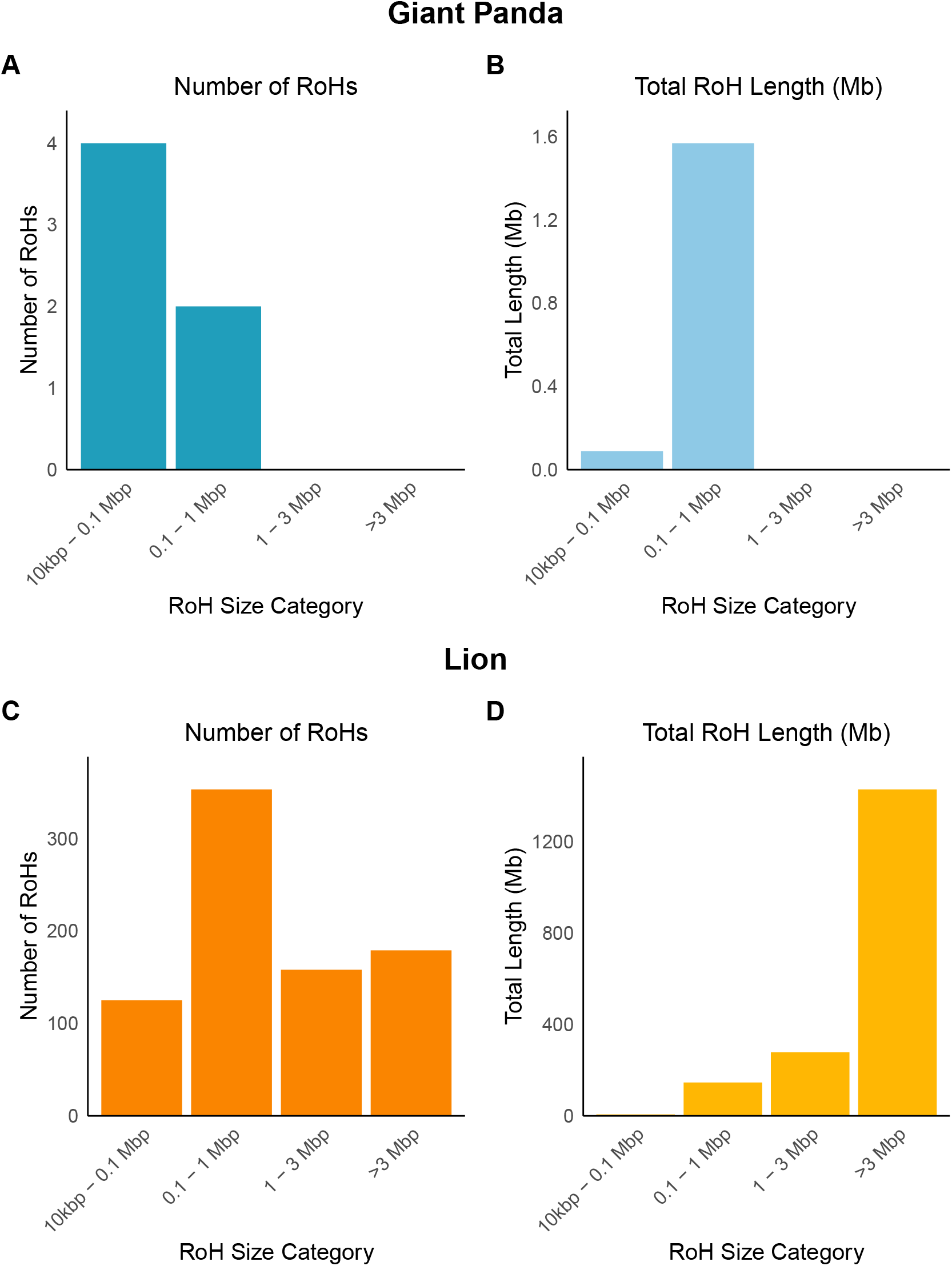
Distribution of Runs of Homozygosity (RoHs) by size in two species. Bar plots of the number of RoHs and the total length of RoHs across four size categories: 10 kbp–0.1 Mbp, 0.1–1 Mbp, 1–3 Mbp, and >3 Mbp. Graphs **A** and **B** show data for the Giant panda, while graphs **C** and **D** show data for the Lion. The left plots (**A** and **C**) show the count of RoHs in each category, while the right plots (**B** and **D**) display the corresponding total genomic length in megabases (Mb).

To illustrate how RoH patterns can differ between species, the distribution in the giant panda genome, characterized by relatively few short RoHs, is shown in Figure 3 A–B. In contrast, the RoH profile of the lion genome, which exhibits a greater number and total length of RoHs, is shown in Figure 3 C-D. These visualizations highlight distinct genomic histories and demographic influences.

### Genome Visualisation

| (Virtual) Machine | Data, Software and Time Requirements |
| --- | --- |
| 28 (V)CPUs<br>64 GB RAM<br>50 GB Storage | **Input:** Reference genome, BAM file and VCF file<br>**Installations:** https://circos.ca/<br>bedtools<br>bcftools<br>mosdepth<br>**Time:** <5h |

Circos plots are widely used in genomics to represent large-scale data in a circular format, allowing for the clear visualization of patterns and connections across entire genomes. This approach is particularly useful for displaying multiple layers of genomic information in a compact and interpretable way. For the SwarmGenomics pipeline, a Circos plot is generated to illustrate key genomic features, including coverage depth, GC content, repeat density, and heterozygosity. These metrics were chosen to provide insights into genome structure and variability. Coverage depth reflects sequencing uniformity, GC content describes sequence composition, repeat density highlights repetitive elements, and heterozygosity indicates genetic diversity.

All metrics are calculated in non-overlapping 1 Mb bins across the genome. Coverage is extracted from BAM files using mosdepth (Pedersen and Quinlan 2018), and repeat density is computed by intersecting repeat-masked regions with predefined genomic bins. GC content is calculated with bedtools nuc (Quinlan 2014), while heterozygosity is determined by using `bcftools view -g het` to filter variant calls from VCF file and counting overlapping variants in each genomic bin.

These results are visualized in a Circos plot with the 20 largest scaffolds of the giant panda genome in Figure 4. GC content and genomic coverage are represented as line plots (purple and red, respectively), while heterozygosity and repeat density are shown as green and blue heatmaps.

**Figure 4:**
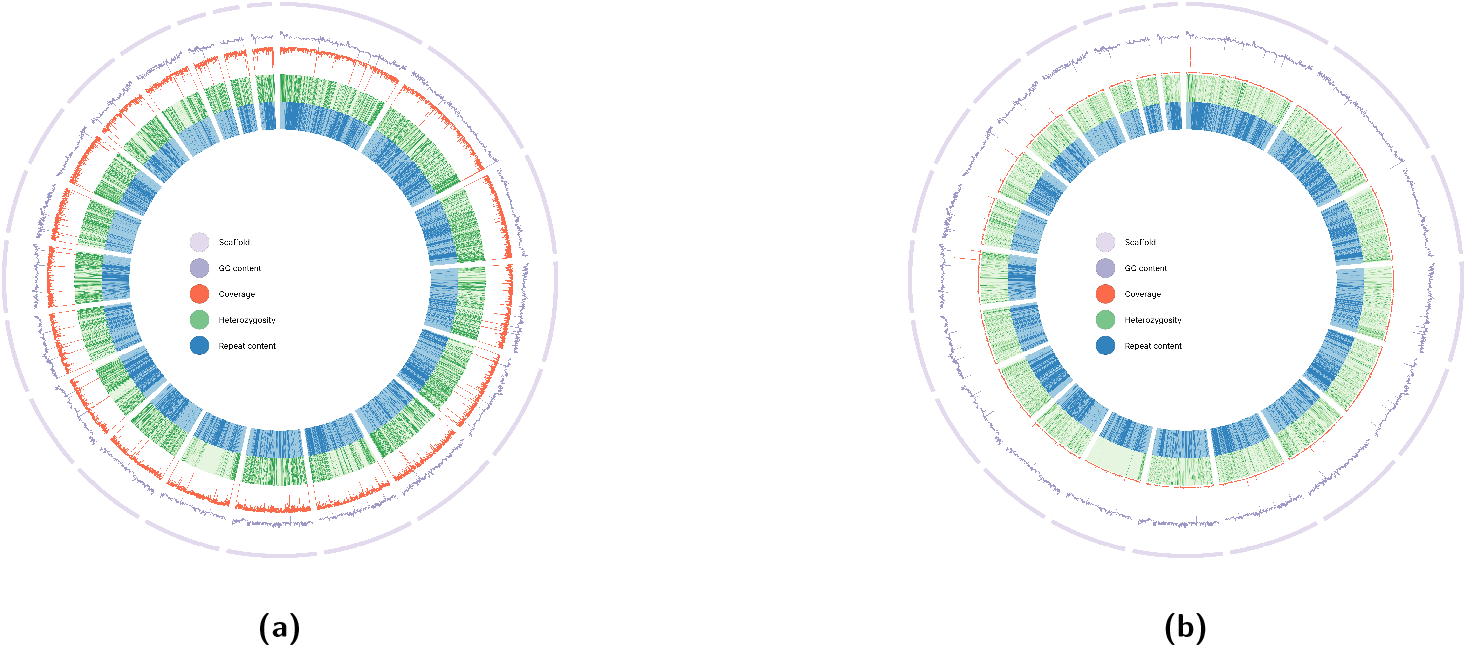
Circos plot displaying genomic features in the 20 largest scaffolds of the giant panda genome: GC content as a purple line, genomic coverage as a red line, heterozygosity as a green heatmap, and repeat density as a blue heatmap. The heterozygosity heatmap ranges from [0–0.002]. **(a)** High coverage genome (50x). **(b)** Low coverage genome (4x).

### PSMC

| (Virtual) Machine | Data, Software and Time Requirements |
| --- | --- |
| 16-28 (V)CPUs<br>32-64 GB RAM<br>100-200 GB storage | **Input:** VCF file<br>**Installations:**<br>PSMC: https://github.com/lh3/psmc.git<br>**Time:** ~ 8h |

The Pairwise Sequentially Markovian Coalescent (PSMC) method provides insights into the historical population sizes of a species, extending back up to millions of years. By analyzing patterns of genetic variation in a single genome, PSMC estimates historical population sizes based on coalescence events. This method is most accurate for periods ranging from 20,000 to 3 million years ago (H. Li and Durbin 2011; Palkopoulou et al. 2015). Estimating recent population sizes can be challenging due to a lack of mutations, while ancient periods are difficult to resolve because of the accumulation of mutations, which introduces noise into the genetic data. By applying PSMC to giant pandas, changes in their population size due to past climatic events, such as the last Ice Age, can be investigated.

PSMC was run following the official guidelines (https://github.com/lh3/psmc) on the diploid consensus sequence `diploid.fq.gz` generated previously. Unless stated otherwise, default settings were used. The **-N** option sets the number of EM iterations (typically 25). The **-t** option specifies the maximum coalescent time (*T*_max_, in units of 2*N*_0_), commonly set to 15. The **-r** option provides the initial estimate of the mutation-to-recombination rate ratio (*θ/ρ*) for the HMM/EM optimization; `-r 5` is a common starting value. The **-p** option defines the pattern of time intervals (free parameters) used to model the TMRCA distribution; a widely used pattern is `“4+25*2+4+6”`. However, this default may occasionally induce spurious recent peaks; splitting the first window (e.g., `“2+2+25*2+4+6”`) mitigates this artifact (Hilgers et al. 2024).

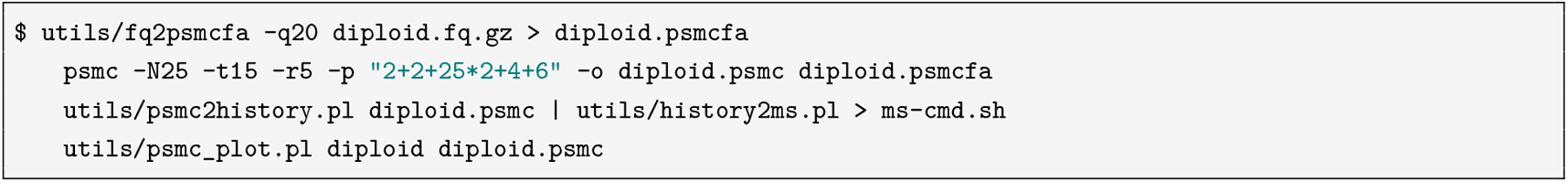

`psmc_plot.pl` from the PSMC program is used for plotting the results, it produces a pdf file with the graph. The script uses the diploid.psmc file, created during PSMC as the input file. To scale the graph according to the species, the user can define generation time (g) and mutation rate (*µ*). Table 2 gives few examples of mutation rates and generation times for different species, which can be used to guide the parameter selection.

**Table 2:** Mutation rates and generation times for various species used in PSMC analyses, as described in the literature.

| Species | Mutation Rate ($\mu$) | Generation Time (years) | Reference |
| --- | --- | --- | --- |
| Human (*Homo sapiens*) | $1.25 \times 10^{-8}$ | 25 | (H. Li and Durbin 2011) |
| Mouse (*Mus musculus*) | $5.4 \times 10^{-9}$ | 0.5 | (Fujiwara et al. 2022) |
| Dog (*Canis lupus familiaris*) | $1 \times 10^{-8}$ | 3 | (Freedman et al. 2014) |
| Great tit (*Parus major*) | $2.0 \times 10^{-9}$ | 2 | (Laine, Gossmann, Schachtschneider, et al. 2016) |
| Chicken (*Gallus domesticus*) | $1.91 \times 10^{-9}$ | 1 | (M.-S. Wang et al. 2021) |

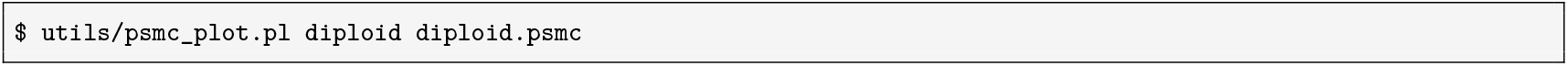

Scaled:

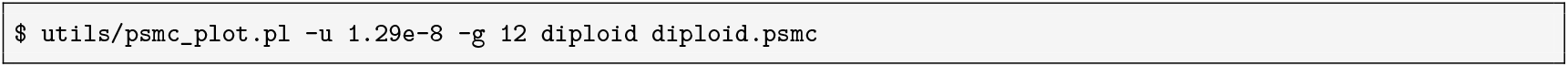

Figure 5 shows the output of `psmc_plot.pl`. The x-axis represents time, with higher values further along the axis, while the y-axis is effective population size. The graph shows a decline in population size at 1,000,000 to 500,000 years ago. This period occurs during the Pleistocene Epoch, which was characterized by repeated glaciations, or ice ages. This could have lead to habitat loss or fragmentation reducing giant panda population sizes.

**Figure 5:**
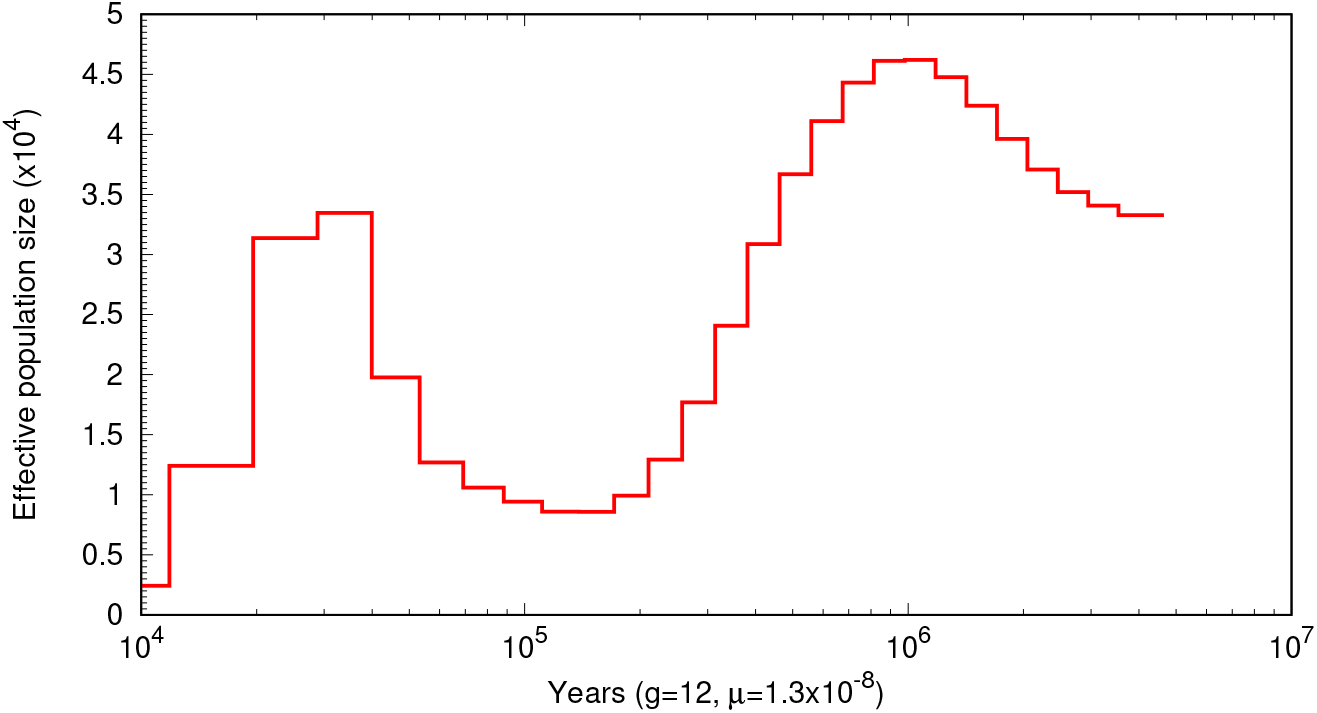
PSMC plot of giant panda populations: Effective population size (x10^4^) is shown on the y-axis, and time (in years) is shown on the x-axis. Parameters used are generation time *g* = 12 and mutation rate *µ* = 1.29 × 10^−8^.

### Mitochondrial genome reconstruction

| (Virtual) Machine | Data, Software and Time Requirements |
| --- | --- |
| 16-28 (V)CPUs<br>32-64 GB RAM<br>100 GB storage | **Input:** FASTQ files<br>**Installations:** GetOrganelle<br>**Time:** <10h |

The study of the mitochondrial genome (mitogenome) is of particular interest due to its unique properties and wideranging applications in genetic research, including genome-wide, epigenome-wide, and phenome-wide association studies (Ferreira and S. Rodriguez 2024). The mitochondrial genome’s high mutation rate, heteroplasmy, copy number variation, maternal inheritance, and lack of recombination make the mitogenome especially valuable in studies of human disease, population structure, evolutionary biology, and DNA barcoding (Y.-H. Lin et al. 2022; Wallace 2010; Song et al. 2022; Jin et al. 2020).

Advances in next-generation sequencing (NGS) technologies have driven the development of numerous specialized tools for organellar genome assembly, particularly for chloroplast and mitochondrial genomes (Dierckxsens et al. 2017). Several assembly tools are described such as GetOrganelle (Jin et al. 2020), NOVOPlasty (Dierckxsens et al. 2017), MitoZ (G. Meng et al. 2019), MITObim (Hahn et al. 2013), SPAdes (Bankevich et al. 2012), MitoFinder (Allio et al. 2020) and Megahit (D. Li et al. 2016), but their assembly qualities vary widely. In a systematic comparison of human mitochondrial genome assembly tools, MitoFlex was the fastest, NOVOPlasty the most memory-efficient for simulated data, GetOrganelle and MitoFlex were best for SNP detection in human WGS data, and MToolBox and NOVOPlasty performed consistently across all sequencing depths (Mahar et al. 2023). Nevertheless, GetOrganelle’s ability to accurately assemble genomes despite complex repeats and highly conserved regions—common challenges in genome assembly—make it a preferred choice for researchers who prioritize accuracy and comprehensive analysis in mitochondrial genome assembly (Jin et al. 2020).

In this pipeline, GetOrganelle is used to assemble the mitogenome from previously generated FASTQ files using the script `“get_organelle_from_reads.py”` with a single line command. This script exploits Bowtie2, BLAST, and SPAdes, as well as the Python libraries NumPy, SciPy, and SymPy as dependencies to filter, align, and assemble reads into contiguous sequences (contigs) representing the organelle genome. The process includes error correction, k-mer analysis, and graph-based assembly. Consequently, the output of GetOrganelle includes the assembled mitochondrial genome in FASTA format, along with detailed assembly statistics and quality metrics. These outputs enable users to perform further analyses, such as annotation of genes, comparative genomics, and evolutionary studies. These results are saved in `DESTINATION_PATH/mt/`, including the assembled mitogenome in FASTA format `(*_mitogenome.fasta)`, the organelle-only assembly graph `(*.selected_graph.gfa)`, log files, and various quality metrics. These outputs are suitable for downstream analyses such as gene annotation, phylogenetic reconstruction, comparative genomics, and population studies (Jin et al. 2020).

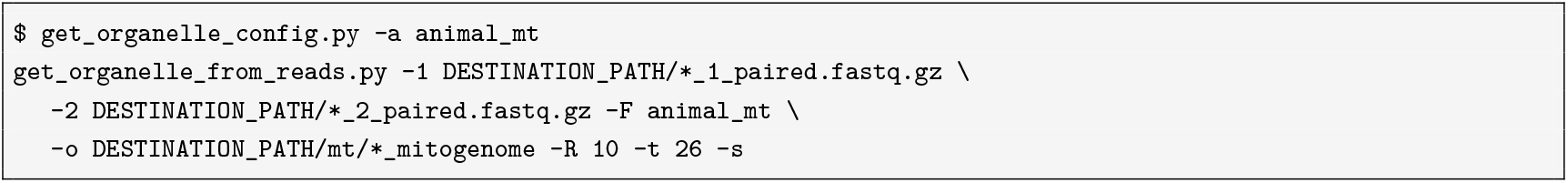

To place the reconstructed mitogenome in a phylogenetic context, a reference database of mitochondrial genome sequences was compiled and formatted using `makeblastdb` (Camacho et al. 2009). The query sequence was renamed for clarity and searched against this database using `BLASTn` (Altschul et al. 1990). The top 10 unique high-scoring matches were selected, combined with the query, and aligned using `MAFFT` (Katoh and Standley 2013). A maximum-likelihood phylogenetic tree was then inferred with `FastTree` based on the resulting alignment (M. N. Price et al. 2010). Full FASTA headers were used as tip labels, with the query sequence highlighted in red as shown in Figure 6a.

**Figure 6:**
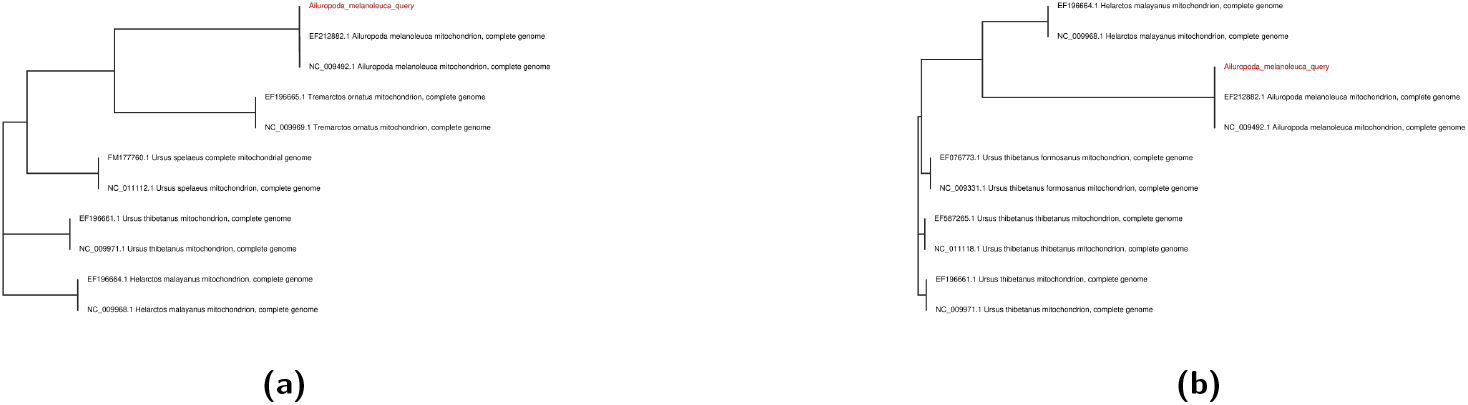
Phylogenetic trees of mitochondrial genomes. (a) Reconstructed giant panda mitochondrial genome highlighted in red among related mitochondrial sequences based on sequence similarity. (b) Reference giant panda mitochondrial genome highlighted in red among related mitochondrial sequences based on sequence similarity.

**Figure 7:**
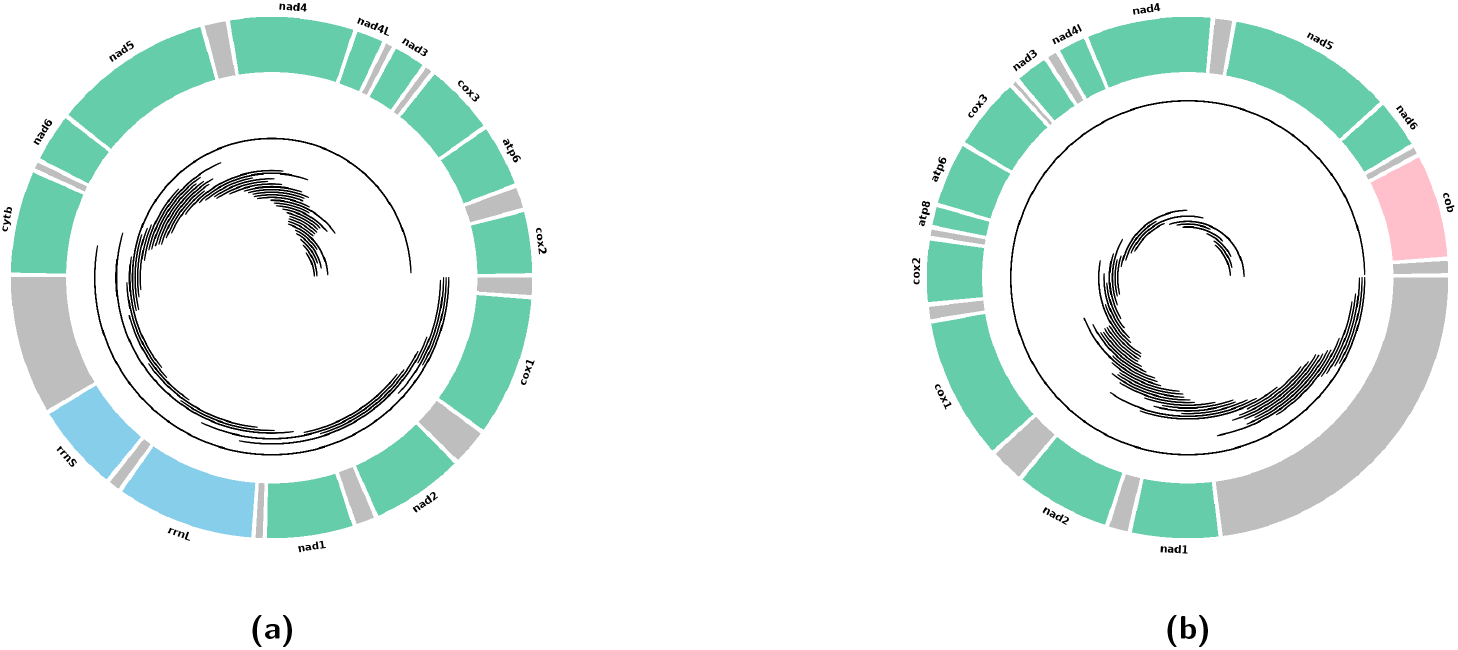
Visualization of NUMT Alignments on the Mitochondrial Genome. The circos plots illustrate the organization of mitochondrial DNA (mtDNA) genes and nuclear mitochondrial DNA segments (NUMTs) aligned to the mtDNA reference. The outermost track displays the annotated mitochondrial genome with genes colour-coded by category: protein-coding genes (green), ribosomal RNA (rRNA) genes (blue), and non-coding regions including introns (gray), other elements are in pink. The inner track shows NUMT alignments as black lines representing the start and end positions of mtDNA regions with homologous sequences in the nuclear genome. **(a)** shows the reassembled mitochondrial genome, while **(b)** depicts the reference mitochondrial genome.

Overall, GetOrganelle represents a significant advancement in the field of organelle genomics, providing researchers with a powerful tool to unlock the genetic secrets of mitochondria and other organelles. Its ability to deliver highquality assemblies from short-read data makes it an invaluable resource for exploring the genetic and functional diversity of these critical cellular components (Jin et al. 2020).

### NUMT identification

| (Virtual) Machine | Data, Software and Time Requirements |
| --- | --- |
| 16-28 (V)CPUs<br>16-32 GB RAM<br>50-100 GB storage | **Input:** Mitogenome FASTA file<br>**Installations:** R, Blast, Bedtools<br>**Time:** <1h |

In this module, NUMT fragments harboured in nuclear genomes are identified. NUMT stands for nuclear mitochondrial DNA, which describes the mitochondrial DNA fragments detected in the nuclear genomes (Lopez et al. 1994). This continuous, ongoing and non-random (W. Wei et al. 2022; Tsuji et al. 2012) integration of mitochondrial DNA to nuclear genome is identified across species and tissues (Hazkani-Covo 2022; Dayama et al. 2020; Liang et al. 2018; Calabrese et al. 2017). The NUMT insertion numbers and sizes vary not only between species but also within populations. In addition, these fragments can introgress from closely related species through hybridization or interbreeding (Bücking et al. 2019; Lammers et al. 2017; Pérez et al. 2017; Baldo et al. 2010), which further fuels the genetic variations in one population. Interestingly, after integration, the NUMT fragments can be preserved in an ancestral state and thus be used for evolutionary event identification and as phylogenetic markers (Uvizl et al. 2024; Lucas et al. 2022; G. Zhang et al. 2021; Miraldo et al. 2012; Soto-Calderón et al. 2014). This shows that NUMTs help unravel the evolutionary and population history. Furthermore, this mutagenic event could be under selection (Vendrami et al. 2022), hinting at the potential function of NUMT.

BLASTN is used to identify NUMTs in the nuclear genome with mitochondrial DNA sequences as queries. First the user has to build the database for BLASTN with nuclear genomes.

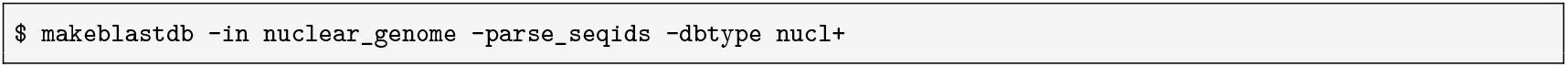

After building the database, the mitochondrial DNA sequences are blasted with the nuclear genome.

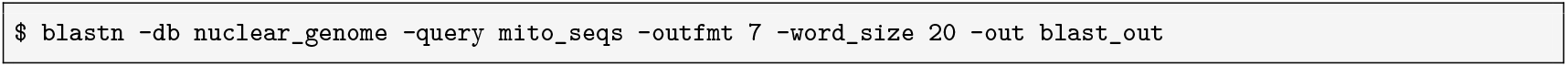

To generate the BLAST results in tabular output with comment lines, which facilitates the creation of the bed file for the next step, the `-word_size` parameter is set to 20. This setting, used in previous research (Lammers et al. 2017; Vendrami et al. 2022), helps identify NUMTs. It should be noted that the `-word_size` parameter can influence the sensitivity and specificity of the BLAST search.

The information is then extracted from the BLAST result using the following command:

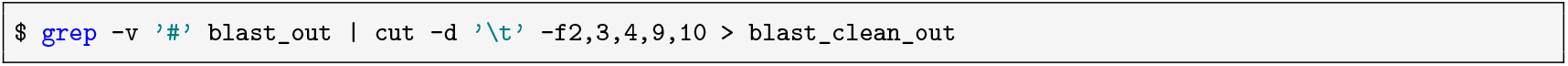

Next, BED files are generated using the provided script `get_bed2.R`. This script not only creates the bed file from the BLAST results but also filters out fragments smaller than 200 bp, to reduce noise in the results.

Finally, `bedtools` is used to generate FASTA files of NUMT sequences from the BED file and the genome file:

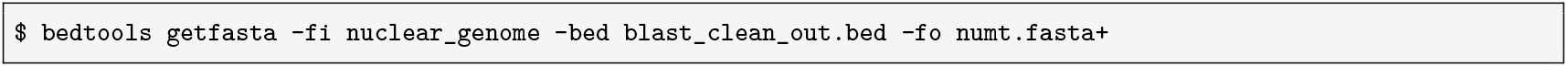

The output file is a FASTA file containing NUMT sequences. Each sequence header includes the scaffold or chromosome name followed by the NUMT position on that scaffold or chromosome. An example pipeline script, `runNUMTidentification2.3.cyc.py`, is available on the GitHub page. The number of NUMTs identified in this genome can be calculated and compared with other species or individuals. The results can be visualised in a circos plot showing the organisation of the mitochondrial genes (mtDNA) and NUMTs aligned Figure **??**. 50 NUMTs were identified in the giant panda, which is consistent with findings in other mammalian species (Y.-C. Chen et al. 2025).

NUMT insertion is an ongoing process that provides valuable information about evolutionary events such as hybridization and interbreeding. After identifying NUMTs, the frequency and extent of this mutagenic event can be examined. The results can also be used for further analyses with other individuals or species, serving as markers for phylogenetic and population genetic studies.

### Repeat analysis

| (Virtual) Machine | Data, Software and Time Requirements |
| --- | --- |
| 12+ (V)CPUs<br>32-64 GB RAM<br>300-500 GB storage | **Input:** Reference genome FASTA file<br>**Installations:** HMMER, RM BLAST, TRF, RepeatMasker, RECON, Ninja, LTR Retriever, Mafft, CD-Hit, Genometools, RepeatModeler2, RepeatScout.<br>**Time:** 24-48h |

Transposable elements (TEs) are ubiquitous genomic components and sources of evolutionary novelty across the eukaryotic tree of life. Indeed, TEs commonly account for the majority of genomic material in both plants and animals, making the study of these self-replicating ‘jumping genes’ an essential component of genomic analyses (Almojil et al. 2021; Slotkin 2018). As TEs evolve within lineages and can ‘jump’ across them, unique TEs often appear within specific clades and can be unexpectedly encountered in any clade. This makes de novo repeat identification necessary for detection of taxon-specific and horizontally transferred TEs (F. Rodriguez and Arkhipova 2022). Annotations of TEs may be of interest for studies on genomic architecture, host-parasite interactions within the genome, and sources of new or laterally transferred genomic material. TE annotations are also commonly considered in population genomic analyses in many species: as TEs commonly result in highly repetitive DNA motifs that interfere with alignment quality and estimates of genomic diversity (Cechova 2020; F. Rodriguez and Arkhipova 2022), TE masking may improve many analyses involving small variants (Patil and Vijay 2021).

This module identifies de novo TE families (Repeats) within a single genome using RepeatModeler2 (Flynn et al. 2020), and annotates a single reference genome for these families, simple repeats, and satellite DNA using RepeatMasker (Tarailo-Graovac and N. Chen 2009). De novo repeat families are identified through the occurrence of similar sequence motifs throughout the genome, eliminating the need for reference TE libraries. Reference sequences are, however, used to assign TE families to classes and subclasses whenever possible, but these are included as DFAM libraries with the RepeatMasker installation and do not require additional inputs from the user. The full script is available on the GitHub page.

Reference-based analysis can also be performed without de novo repeat annotation. This can be done very quickly and with minimal prerequisites. It is, however, most informative when studying narrow groups within well-annotated clades: Applying it to mammals (‘mammalia’) and birds (‘aves’) will provide fast and informative insights into the repeat landscapes of most species within these classes. Reference-based methods on poorly studied taxa may not yield meaningful results, and analyses with very broad categories of reference data (e.g. eukaryota) will likely exceed the run times required for de novo annotation.

Running this analysis requires only a single reference genome as input. The user may also wish to set the number of parallel threads to be used. This module will produce a FASTA file of de novo repeat families (“…-familes.fasta”) broken into subfamilies with greater than `80%` identity to one another. It will also produce a file of repeat annotations on the reference genome (“…fna.out”), a summary table of repeats annotated within the reference genome (“…tbl”), an interactive summary graph of the TE landscape as the genomic proportion of repeats by divergence from consensus (“…html”), and a repeat masked reference genome for downstream analyses (“…fna.masked”). This will provide the user with the necessary files for a general analysis on TEs within an assembly, and allow for the exclusion of TEs from associated population genomic analyses as required.

The repeat landscape of the giant panda genome is visualised in Figure 8 with the fraction of different repeat families in the genome as a pie chart. Repeats make up `31%` of the genome with LINE families being most abundant. The y-axis of the repeat landscape shows the percentage of the genome that each repeat family makes, with the age of the repeats represented as the Kimura substitution level on the x-axis. In the giant panda genome, it shows a recent expansion of the LINE1 family, possibly indicating active repeats.

**Figure 8:**
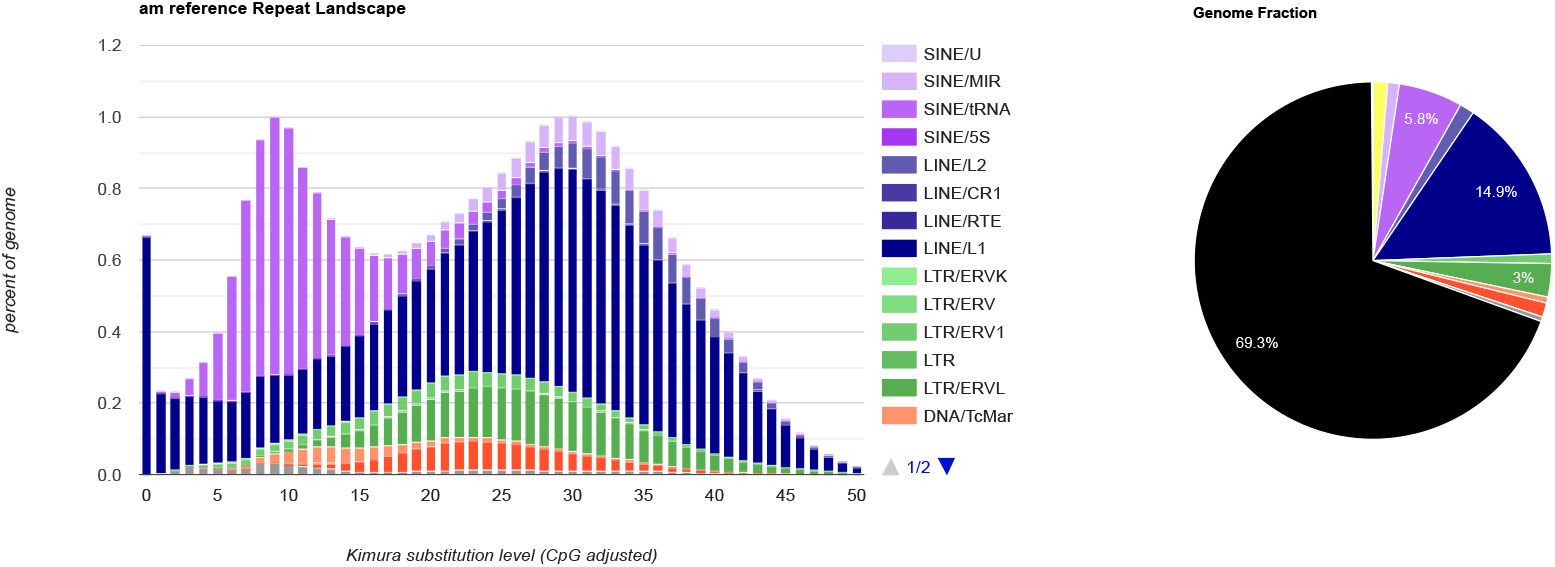
Repeat landscape of the giant panda genome: The percentage of repeat families in the genome is shown on the y-axis, while the Kimura distance is plotted on the x-axis. A pie chart representing the percentage distribution of repeat families in the genome is also included.

### Unmapped reads classification

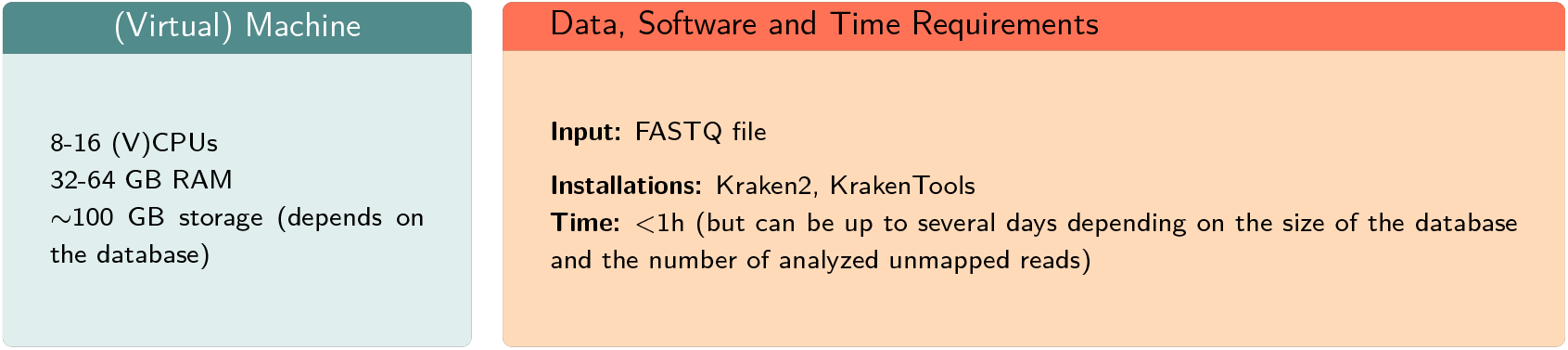

| (Virtual) Machine | Data, Software and Time Requirements |
| --- | --- |
| 8-16 (V)CPUs<br>32-64 GB RAM<br>~100 GB storage (depends on the database) | **Input:** FASTQ file<br>**Installations:** Kraken2, KrakenTools<br>**Time:** <1h (but can be up to several days depending on the size of the database and the number of analyzed unmapped reads) |

An important source for UR may be microbes that are not contaminations but actually belong to the microbiome of the sampled organism. Bacteria are often important parts of the microbiome and may may persist in body parts once considered sterile, such as eggs, blood, lung and bladder (D’Aquila et al. 2021; O’Dwyer et al. 2016; Nyholm 2020; Trevelline et al. 2018). Viruses can also be studied by analysing UR (Moustafa et al. 2017). Such UR of viral origin might come from viruses that historically integrated into the host genome of a population, viruses that are integrated into the genome of only the sampled individual host, or viruses from an acute infection at the time point of sampling. Finally, genomic variation of the host can be a source of UR. For example, reads with complex structural variations, like large indels, inversions and translocations may fail to map to the reference, if reads are mapped to a reference genome, which does not include such structural variations. This may happen when the reference individual and the sampled individual are from populations with divergent structural variants (Takayama et al. 2021; K. H. Wong, Levy-Sakin, et al. 2018; K. H. Wong, Ma, et al. 2020). Altogether, there are multiple sources of variation in UR, which differ in their biological relevance, and their study can facilitate the understanding of genomic architecture and host-symbiont relationships.

WGS studies typically begin with assembling the genome of the organism of interest. The assembled genome then serves as a reference for subsequent whole-genome re-sequencing, in which sequenced reads from a newly generated sample are mapped to the reference genome. In such cases, a certain part of the produced reads cannot be mapped to the reference genome. These unmapped reads (UR) may result from methodological artifacts, sequencing errors, and they are typically ignored and excluded from further analyses (Chrisman et al. 2022). However, many of them may still contain interesting biological information (Chrisman et al. 2022). For example, they were used to identify biotype-specific sequences in the aphid *Acyrthosiphon pisum* and its endosymbionts (Gouin et al. 2015), and to detect the blood parasites *Plasmodium* and *Tryptpanosoma* in RNA samples of the Great Tit *Parus major* (Laine, Gossmann, Van Oers, et al. 2019).

In this module, Kraken 2 enables taxonomic classification of metagenomic reads (Wood et al. 2019). It compares read *k*-mers to a database of *k*-mers of known origin. For the analysis, Kraken 2 requires a FASTA file with the unmapped reads to be classified and the database to compare these reads to. For the following example, Kraken 2 and Krakentools have been pre-installed, e.g. in a miniconda environment called “kraken”, which can be activated as follows.

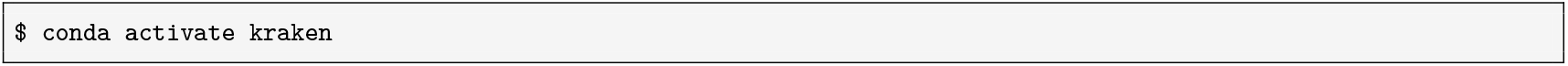

First, the user must download and build a database containing the *l* -mers and the taxonomic information. One can (i) download a pre-built database from https://benlangmead.github.io/aws-indexes/k2, (ii) use the `kraken2-build --standard` command, or (iii) build one’s own custom database with all the taxa that should be included. For simplicity, the use of a pre-built database will be described here. However, it should be noted that building a custom database may be necessary to achieve the best and most meaningful results.

Here, we used the PlusPFP-16 database, a database that adds sequences of protozoa, fungi, and plants to those in the standard database and is capped at 16 GB. PlusPFP-16 is downloaded (by clicking on the ‘.tar.gz’ file in the relevant row), and the tarball can then be extracted.

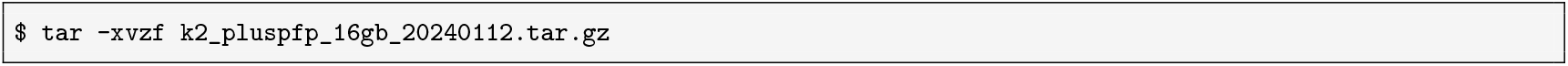

For classifying the unmapped reads from a whole-genome sequencing project, the `kraken2` command is used. The placeholder `$DBPATH` should be replaced with the path where the database is saved, along with the name assigned to the database. The number of threads used by Kraken 2 can also be specified by replacing `$THREADNUM` with the desired number of threads. The flags `--classified-out` and `--unclassified-out` are optional.

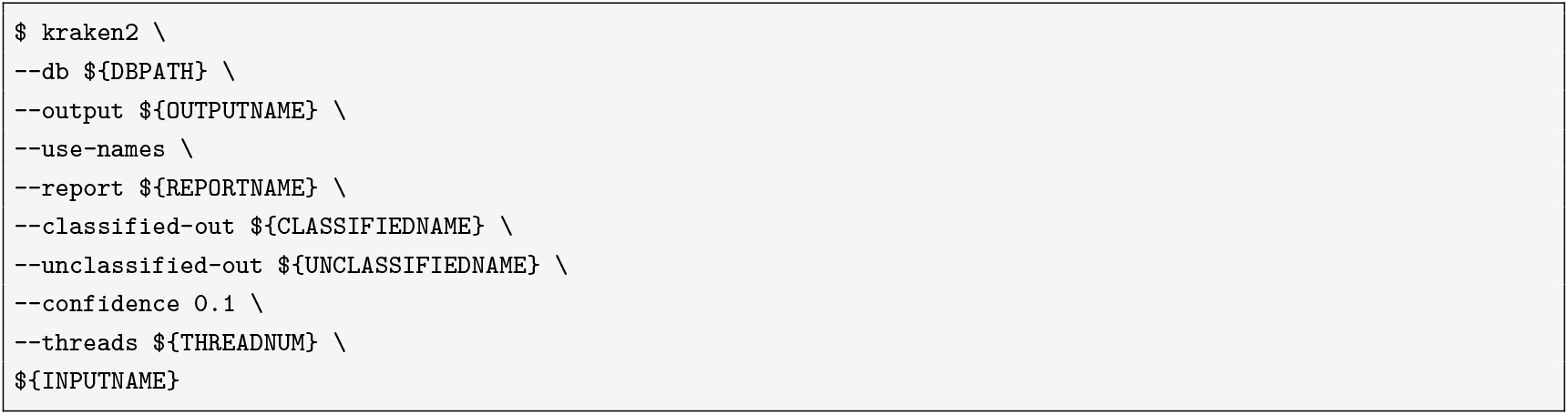

Kraken 2 requires sufficient free memory to load the database into RAM, i.e. at least 16GB free RAM in this case. This command may take some time to complete and will generate a classification output file (.kraken) and a taxonomic summary (.kreport). The output file (with a `.kraken` extension) contains a list of all reads and their classifications. The report (with a `.kreport` extension) is particularly useful, as it shows the number of reads classified under each taxon.

At this point, the script `kreport2krona.py` from KrakenTools can be used to create a visualization of the results.

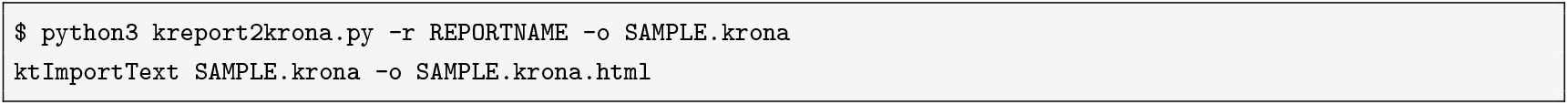

The final results show that from the `1%` classified unmapped reads, nearly all came from humans, which suggests that during the sample collection or laboratory work the giant panda sample was contaminated with human DNA. `0,09%` of the classified unmapped reads had bacterial origin (Figure 9). In addition, a small fraction has come from viruses. Still, `99%` of the unmapped reads were unclassified, suggesting that building a larger custom database might be beneficial.

**Figure 9:**
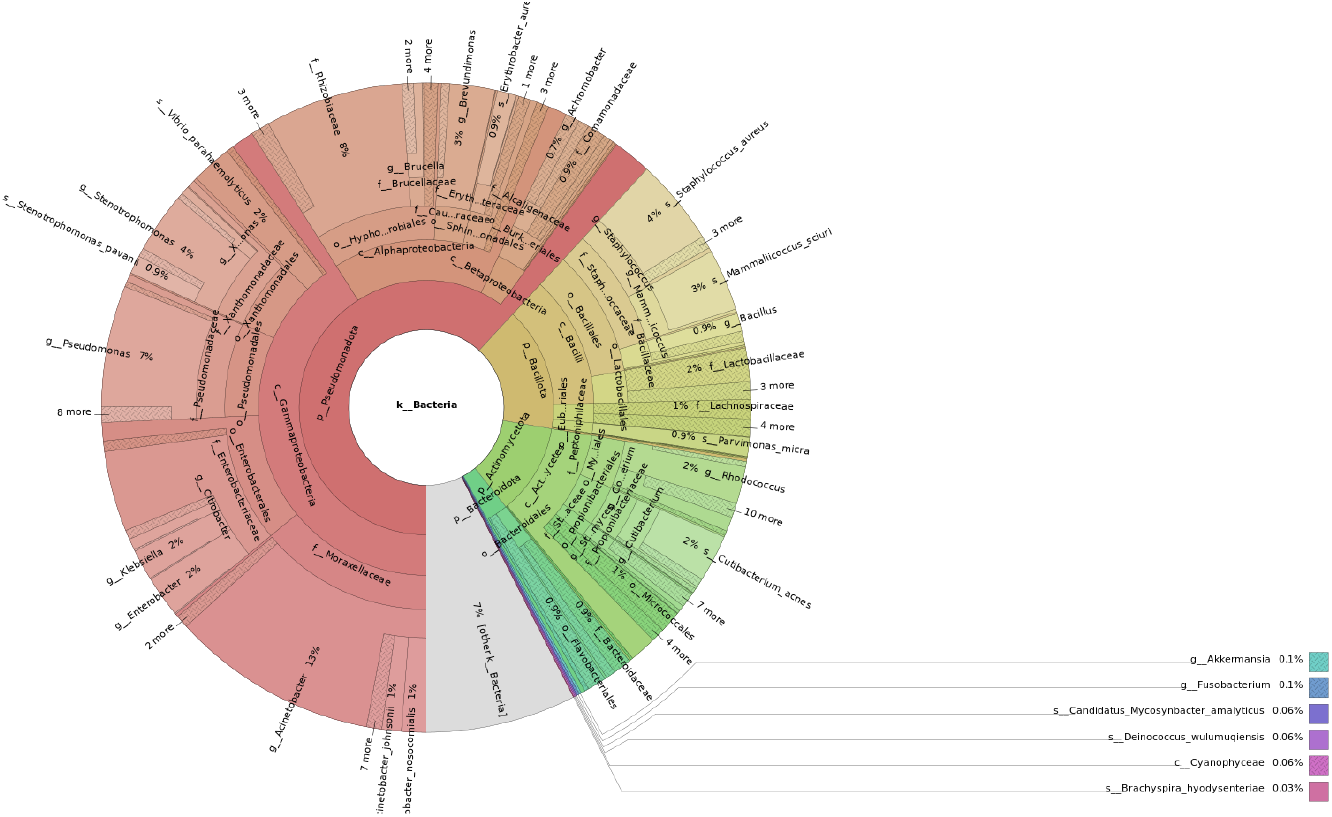
Visualization of the classified bacterial reads from the genomic sample of the giant panda: The graph shows the proportion of classified bacterial reads in the unmapped data.

## Comparative genomics

Having access to existing genomic data greatly enhances the interpretability and context of newly analysed genomes. As users process their data through SwarmGenomics, they can directly compare key metrics, such as heterozygosity, repeat content, or inbreeding levels—to those from a growing database of diverse species.

To illustrate the utility of comparative genomic analyses, several summary plots are presented. Scaffold continuity is compared across species by showing the proportion of scaffolds above and below 5 Mb (Figure 10a). Heterozygosity levels are examined in relation to body mass, obtained from MammalBase (The MammalBase Community 2022) and Avibase (Lepage et al. 2014), providing insight into potential life-history patterns (Figure 10d). Runs of homozygosity are visualized to highlight extreme cases, such as the Indian lion, known for its inbred genome (Figure 10c) (De Manuel et al. 2020). The relationship between NUMTs and repeat content is explored across genomes (Figure 10b). Lastly, the most abundant bacterial phyla detected from unmapped reads are summarized, providing a snapshot of the potential microbiome or environmental signals in the raw sequencing data (Figure 10e)

**Figure 10:**
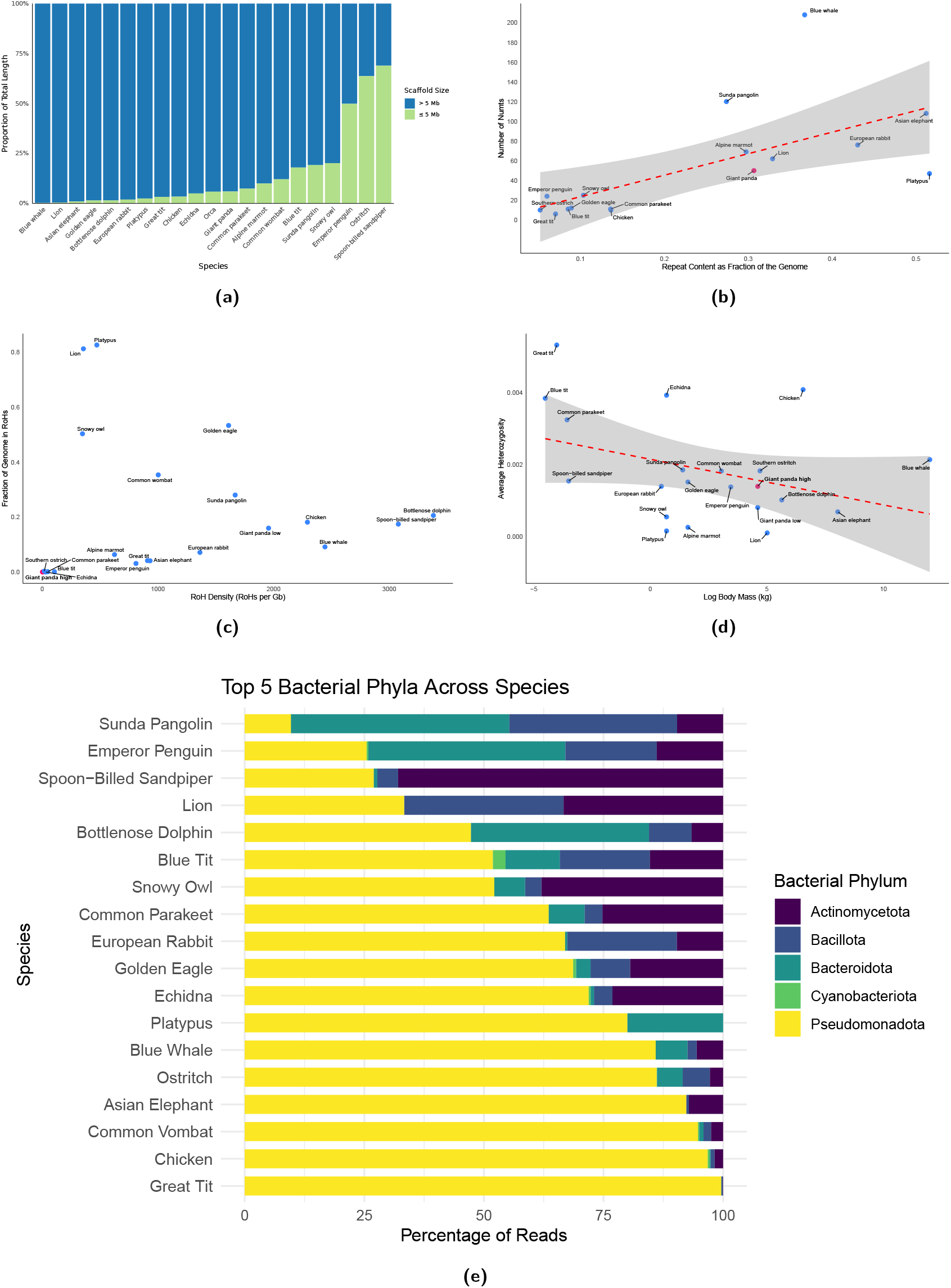
Comparative genomics and associated analyses across studied species. (a) Proportion of scaffolds below and above 5 Mb across the genomes. (b) Correlation between NUMTs and repeat elements across species. (c) Comparison of Runs of Homozygosity (ROH) across species. (d) Correlation between heterozygosity and body mass. (e) Top 5 bacterial phyla across the species.

## Discussion

The SwarmGenomics pipeline was developed to simplify whole-genome assembly and streamline downstream analyses, providing insights into species-level genomic diversity and evolutionary history. The pipeline comprises seven modular components: heterozygosity, runs of homozygosity, PSMC modeling, unmapped reads, repeat annotation, mitochondrial genome assembly, and NUMT analysis. Together, these modules demonstrate how meaningful genomic characteristics can be extracted from a single whole-genome sequence, even for non-model species.

Using a single giant panda genome, we uncovered valuable insights into the genetic health and history of the species. The heterozygosity analysis across the largest scaffolds revealed moderate genetic diversity, although values varied by scaffold. This aligns with earlier studies using microsatellites, which have also shown variability in genetic diversity across panda populations (Lu et al. 2001; Shan et al. 2014; W. Li et al. 2023). The RoH analysis revealed few runs of homozygosity longer than 10 kb, suggesting an absence of recent inbreeding in the sampled individual. In contrast, application of the same analysis to a lion genome revealed numerous long RoHs, reflecting a different inbreeding history. The PSMC modeling indicated a major decline in effective population size between 500,000 and 1 million years ago, corresponding to two major Pleistocene glaciations in China, consistent with findings in (Zhao et al. 2013). Additionally, we identified 50 NUMTs, which could potentially be studied further to investigate historical hybridization or introgression events. The repeat annotation revealed a recent expansion of the LINE1 transposable elements, possibly indicating ongoing activity. TEs overall were found to constitute nearly one-third of the genome, consistent with previous estimates of approximately `37%` (R. Li et al. 2010). The unmapped reads module identified signs of contamination by human DNA, as well as sequences of bacterial origin, with a large fraction attributed to the Pseudomonadota phylum. Given that only `∼1%` of unmapped reads could be classified, a larger, custom database would likely improve resolution. Altogether, the analysis illustrates how a single genome can reveal insights into genetic diversity, demographic history, symbiont presence, and evolutionary processes relevant to species conservation and biology.

### Pipeline Features and Future Development

A key strength of SwarmGenomics is its modular design, which enables users to select only the analyses most relevant to their study system and research questions. At present, the pipeline assumes access to a pre-assembled reference genome, but future releases will include a module for reference genome assembly, broadening applicability to a wider range of species.

Additional planned expansions include modules for population genetic inference, evolutionary genomics, and selection analysis, such as evaluating coding region heterozygosity and SNPeffect. We also aim to support multiple sequencing platforms, including PacBio and Oxford Nanopore, which will allow the incorporation of epigenomic data in future versions of the pipeline.

### Collaborative Development and Educational Use

The long-term vision of SwarmGenomics is to function as both a bioinformatics teaching platform and a collaborative research resource. The project’s GitHub repository provides all materials needed to run analyses, with user-friendly documentation designed to support students and researchers new to genomics. It is freely available for use in university courses and self-guided learning.

A dedicated SwarmGenomics website is currently under development and will serve as a central database for user-generated results. Users will be able to contribute intermediate and final output files, which will be made publicly accessible and downloadable, facilitating open data sharing.

“SwarmGenomics” refers to the project’s emphasis on collaboration, where multiple researchers contribute genomic datasets and analyses to a common platform. This open-access model enhances the reuse of data and allows comparative studies across diverse taxa. By enabling shared contributions and reproducible workflows, SwarmGenomics aims to strengthen collaborative research in evolutionary and conservation genomics.

## Conclusion

In conclusion, SwarmGenomics offers a comprehensive and user-friendly pipeline that integrates whole-genome assembly with advanced genomic analyses. By providing insights into population genetics and evolutionary history, it serves as a valuable tool for both research and education. Planned developments, including modules for reference genome assembly and expanded analytical capabilities, will further enhance its utility across diverse taxa. We invite the community to use, adapt, and contribute to SwarmGenomics, promoting a shared platform for advancing genomic research and discovery.

## Software availability statement

SwarmGenomics is available on GitHub at https://github.com/AureKylmanen/Swarmgenomics

## Author contribution

TIG designed the study with input from all authors. AK implemented the pipeline and performed the analyses with help from all other authors. AK drafted the manuscript with input from all authors. TIG provided supervision, critical feedback, and manuscript revisions. All authors read and approved the final manuscript.

## Acknowledgments

This work was supported by funding from the European Research Council (ERC) under the European Union’s Horizon 2020 research and innovation programme, grant agreement No. 947636. This work was supported by the de.NBI Cloud within the German Network for Bioinformatics Infrastructure (de.NBI) and ELIXIR-DE (Forschungszentrum Jülich and W-de.NBI-001, W-de.NBI-004, W-de.NBI-008, W-de.NBI-010, W-de.NBI-013, W-de.NBI-014, W-de.NBI-016, W-de.NBI-022). We would like to thank Ina Karabay for providing a baseline script for genome pre-processing. We also appreciate the students of the SwarmGenomics course at TU Dortmund University for their participation, which has allowed us to refine and enhance the course further. Moreover, we wish to express gratitude to other members of Computational Systems Biology group for helpful feedback.

## Glossary

Diploid Organism: An organism that has two sets of chromosomes, one from each parent.
Genome Assembly: The process of piecing together short DNA sequences to reconstruct the full genome of an organism.
Heterozygosity: A measure of genetic variation indicating the presence of different alleles at a gene locus within an individual or population.
Runs of Homozygosity (RoH): Contiguous stretches of DNA in the genome where the alleles at all loci are homozygous — indicating that identical copies of genetic material were inherited from both parents across this region.
PSMC (Pairwise Sequentially Markovian Coalescent): A computational method used to infer historical population sizes from a single diploid genome.
Unmapped Reads: Sequences in a genomic dataset that cannot be aligned to a reference genome, potentially indicating novel or uncharacterized DNA.
Repeat Analysis: The study of repetitive DNA sequences in the genome, which can have structural and regulatory functions.
Mitochondrial Genome Assembly: Reconstruction of the complete mitochondrial DNA sequence from whole-genome sequencing data.
NUMTs (Nuclear Mitochondrial DNA Segments): Fragments of mitochondrial DNA that have been incorporated into the nuclear genome.
Effective Population Size: The number of individuals that effectively participates in producing the next generation. It reflects the genetic diversity of a population, which can be different from the actual number of individuals in the population.
Reference Genome: A high-quality, complete genome sequence used as a standard for mapping and comparing DNA sequences from other individuals or species.
PacBio/Nanopore Sequencing: Advanced sequencing technologies that produce long-read sequences, useful for assembling complex genomes and studying epigenetics.
Variant Calling: The process of identifying genetic variants, such as SNPs or indels, by comparing sequencing data to a reference genome.

## FAQs

### Q1: Can I use the pipeline for [X] species?

SwarmGenomics is currently optimized for diploid species. While it may run on polyploid organisms, this can lead to inaccurate results due to incorrect assumptions about genome structure and heterozygosity.

### Q2: What types of input data does SwarmGenomics accept?

To run the full pipeline, SwarmGenomics requires a reference genome in FASTA format, ideally high quality and species specific, and raw sequencing reads from a single individual in FASTQ or SRA format. These can come from the same individual or from different individuals of the same or a closely related species.

While both inputs are needed to complete the full workflow, some modules can be run independently with alternative input formats (see Table 1 for details).

### Q3: Is there a recommended sequencing depth or genome size limit for using the pipeline effectively?

SwarmGenomics works best with sequencing coverage around 20–30× for the diploid individual to ensure reliable results. It can be run on lower coverage data, but results may be less accurate and less reliable. There is no strict genome size limit, though larger genomes will require more computational resources.

### Q4: Why is a virtual machine recommended?

Many SwarmGenomics modules require substantial computational resources that may exceed the capacity of a typical personal computer, so using a VM on a high-performance server or cluster helps manage these demands efficiently.

### Q5: Do I need internet access throughout the pipeline run? An internet connection is required to connect to the virtual machine and during software installations. However, smaller operations may be run locally without an internet connection

### Q6: How do I install the required software for SwarmGenomics? We recommend following the official installation instructions available in the SwarmGenomics GitHub repository: https://github.com/AureKylmanen/Swarmgenomics/

The repository includes a step-by-step installation guide and optional setup scripts to help you install all required dependencies using `conda`, download additional tools like the SRA Toolkit and BCFtools, and configure your environment correctly (including for de.NBI VMs).

For best results, please read the instructions carefully and ensure that you have the appropriate system permissions, especially when installing on shared or cluster environments.

### Q7: How to download the reference genome assembly?

To download a reference genome, we recommend using the NCBI Genome Data page: https://www.ncbi.nlm.nih.gov/datasets/genome/.

1. Search for your species of interest.
2. On the species page, select the **FTP** tab to view the available files.
3. Look for the file ending in `.genomic.fna.gz` — this is the genome assembly file.
4. Copy the link to this file and use the `wget` command in your terminal to download it. For example:

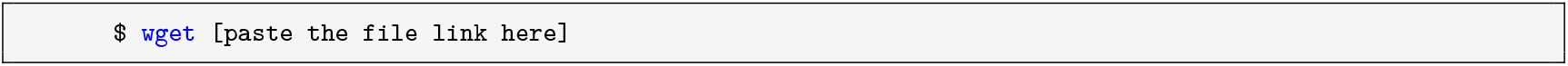

### Q8: How to download raw sequencing reads (in FASTQ or SRA format?)

To download high-quality raw sequencing reads for whole-genome assembly, we recommend using the NCBI Sequence Read Archive (SRA): https://www.ncbi.nlm.nih.gov/sra.

1. Use the **Advanced Search** feature to filter datasets. Recommended search filters include:
  - Platform: Illumina
  - Source: DNA
  - Strategy: Genome
  - Layout: PAIRED
  - File Type: FASTQ or SRA
2. Choose a dataset with a large number of reads (e.g., ≥10 GB compressed)
3. Once you’ve selected a suitable dataset (e.g., with an accession like `SRR12345678)`, you can download it using `sra-tools`.

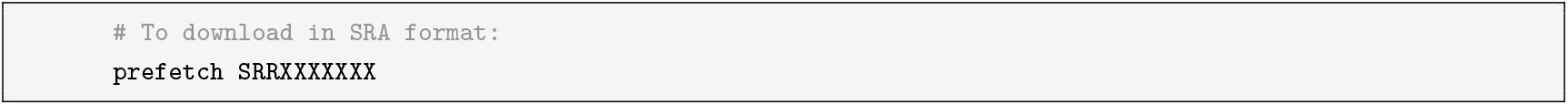

Make sure `sra-tools` is installed and properly configured before running these commands.

### Q9: What does the “No such file or directory” error mean, and how can I fix it?

The “No such file or directory” error in bash means the system can’t find the file or folder you’re trying to access. This often happens because of typos, incorrect file paths, or because the file doesn’t exist in the current directory.

To fix it, check that the filename and path are correct (remember Linux is case-sensitive), use ls to list files in the directory, and verify your current location with pwd.

### Q: Still have questions?

If you have any additional questions or need further assistance, feel free to contact us. We’re happy to help!

## Notes

https://github.com/AureKylmanen/Swarmgenomics

### Competing Interest Statement

The authors have declared no competing interest.

